# The *Arabidopsis thaliana* core splicing factor PORCUPINE/SmE1 requires intron-mediated expression

**DOI:** 10.1101/2024.07.02.601721

**Authors:** Varvara Dikaya, Nelson Rojas-Murcia, Ruben M. Benstein, Wolf L. Eiserhardt, Markus Schmid

**Affiliations:** Umeå Plant Science Centre, Department of Plant Physiology, Umeå University, SE-901 87 Umeå, Sweden; Department of Plant Biology, Linnean Center for Plant Biology, Swedish University of Agricultural Sciences, S-75007 Uppsala, Sweden; Department of Biology, Aarhus University, Ny Munkegade 114-116, 8000 Aarhus C, Denmark; Royal Botanic Gardens, Kew, Richmond, Surrey, TW9 3AE, UK

**Keywords:** *Arabidopsis thaliana*, alternative RNA splicing, temperature signaling, SmE, PORCUPINE (PCP)

## Abstract

Plants are prone to genome duplications and tend to preserve multiple gene copies. This is also the case for the genes encoding the Sm proteins of *Arabidopsis thaliana* (L). The Sm proteins are best known for their roles in RNA processing such as pre-mRNA splicing and nonsense-mediated mRNA decay. In this study, we have taken a closer look at the phylogeny and differential regulation of the SmE-coding genes found in *A. thaliana*, *PCP/SmE1*, best known for its cold-sensitive phenotype, and its paralog, *PCPL/SmE2*. The phylogeny of the *PCP* homologs in the green lineage shows that *SmE* duplications happened multiple times independently in different plant clades and that the duplication that gave rise to *PCP* and *PCPL* occurred only in the Brassicaceae family. Our analysis revealed that *A. thaliana* PCP and PCPL proteins, which only differ in two amino acids, exhibit a very high level of functional conservation and are able to perform the same function in the cell. However, our results indicate that *PCP* is the prevailing copy of the two *SmE* genes in *A. thaliana* as it is more highly expressed and that the main difference between *PCP* and *PCPL* resides in their transcriptional regulation, which is strongly linked to intronic sequences.

## Introduction

Rooted in place, plants cannot escape changes in their environment. Instead, they constantly monitor their surroundings and adjust their growth and physiology to ensure survival. Daily and seasonal temperature fluctuations in particular are important environmental cues that, together with other external signals, inform plants about their surroundings. The actual responses differ depending on the temperature the plant experiences. For example, moderately elevated temperature triggers a reaction known as thermomorphogenesis whereas severe heat stress can stop plant growth altogether and ultimately result in the death of the plant (Quint et al. 2016; Hasanuzzaman et al. 2013). Similarly, cold is usually divided into low permissive temperatures that only mildly affect plant development, chilling that induces a cold and general stress response, and temperatures below 0°C, or freezing, that might cause ice formation and severe damage inside the plant cell. Importantly, exposure to chilling conditions can prime and prepare plants for a further temperature drop (Chinnusamy et al. 2007; Baier et al. 2019). Depending on their habitat, plants may face either only chilling, freezing, or both.

Perception of temperature in plants is mediated by a combination of multiple signaling pathways rather than a single dedicated sensor. Initially, temperature modulates the physical properties of cellular components such as the rigidity of membranes, enzymatic activity, conformational changes of protein and nucleic acid structures, (dis-) assembly of the cytoskeleton, and changes in the viscosity of solutions (Graham and Patterson 1982; Guy 1990; Örvar et al. 2000; Ruelland and Zachowski 2010). These primary physical responses initiate signals via multiple pathways that work simultaneously: MAPK cascades, Ca2+ influx into the cell, and rearrangement of transcription and translation (Miura and Furumoto 2013). Many players of cold signaling are not specific to temperature responses but also participate in other signal pathways such as light, osmotic stress, drought, and general stress response (Lamers et al. 2020; Manasa S et al. 2022). The ultimate result of the signal transmission is an adjustment of the gene regulation, metabolism, overall developmental program of the plant, and its acclimatization to the current growth conditions. Not surprisingly given the scope and importance of temperature regulation, multiple mechanisms, including regulation at the epigenetic level, transcription, translation, and post-translational modifications, contribute to this process (Manasa S et al. 2022).

RNA splicing (Berget et al. 1977;Gilbert 1978), the process during which non-coding introns are removed from the pre-mRNA and exons are joined, has emerged as a key regulatory mechanism that enables plants to respond to and adjust their physiology, growth, and development to their surroundings (Syed et al. 2012;Staiger and Brown 2013; Shang et al. 2017; Liu et al. 2022). Conceptually, RNA splicing is often divided into constitutive splicing (CS), during which a prevalent RNA isoform is formed, and alternative splicing (AS), which enables the organism to produce multiple distinct mature mRNAs from one pre-mRNA (Kelemen et al. 2013; Reddy et al. 2013). In *A. thaliana* about 60% of genes undergo AS (Filichkin et al. 2010;Marquez et al. 2012; Zhang et al. 2022).

AS has been shown to play a role in establishing organ identity, regulating transitions between developmental stages, and contributing to the response of plants to various environmental stimuli (Szakonyi and Duque 2018; Laloum et al. 2018; Tognacca et al. 2023). For example, it has been shown that the number and type of AS events in the cell change significantly in response to cold and approximately 27% of the cold-responsive genes showed differential AS in *A. thaliana* (Calixto et al. 2018). However, even though multiple observations demonstrate the importance of RNA splicing in coordinating a plant’s response and acclimation to temperature (Jiang et al. 2017; Li et al. 2020a; Zhong et al. 2024), the contribution of individual splicing factors and their connection to canonical temperature signaling pathways is largely unknown (Dikaya et al. 2021).

A gene that has recently been shown to be important for the proper growth and development of *A. thaliana*, particularly under cool ambient temperatures and in cold conditions, is *PORCUPINE* (*PCP*; *SmE1*; At2g18740) (Capovilla et al. 2018; Huertas et al. 2019; Wang et al. 2022). *PCP* encodes an E-subunit of the evolutionary conserved Sm-ring that together with U-rich small nuclear RNAs forms the core of the U1, U2, U4, and U5 small nuclear ribonucleoproteins (snRNPs) (Khusial et al. 2005; Will and Lührmann 2011). All members of the Sm-ring are highly conserved in their structure and possess an N-terminal α-helix followed by a five-stranded antiparallel β-sheet which forms characteristic Sm1 and Sm2 motifs (Hermann et al. 1995; Kambach et al. 1999; Khusial et al. 2005). Structural analyses of the human Sm-ring revealed that the ring-like structure is formed by the interaction between the β4 strand of one Sm-protein and the β5 strand of its counter-clockwise neighbor (Weber et al. 2010). The Sm-site that binds snRNA inside the central opening of the Sm-ring is formed by residues that are positioned in the loops L3 (between strands β2 and β3) and L5 (between β4 and β5) and belong to the Sm1 and Sm2 motif, respectively (Weber et al. 2010). Together with other proteins, snRNPs form the core of the spliceosome, the molecular machinery that catalyzes the splicing reaction (Will and Lührmann 2011). An intriguing feature of *A. thaliana* plants deficient in *PORCUPINE* is that these mutants appear macroscopically like wild-type plants at 23°C but display severe developmental phenotypes at moderately cool temperatures of 16°C including significant aberrations in shoot- and root apical meristem development, delayed flowering time, and male sterility (Capovilla et al. 2018). More recently it has been shown, however, that the *pcp-1* mutant exhibits subtle phenotypes at the cellular level, for example in the root meristem, when grown at a temperature of 23 °C (El Arbi et al. 2024). These findings demonstrate that the *PCP* gene is important for normal development across the entire temperature range but that mutant phenotypes manifest mostly at cool temperatures.

Sm genes have been duplicated in plants, and often two or more copies of each subunit are found in a given species (Chen and Cao 2014; Gao et al. 2021). This is also the case in *A. thaliana* where each of the seven subunits that form the Sm-ring has been duplicated (Cao et al. 2011). The coding sequences of *A. thaliana PCP* (SmE1) and its paralog *PCP-LIKE* (PCPL; SmE2) are highly similar and the two proteins differ in only two amino acids (Capovilla et al. 2018). The biological function of PCPL has, however, not yet been investigated in detail.

Here we have conducted a detailed phylogenetic analysis of SmE proteins in the green plants (Viridiplantae). It revealed that duplication of SmE likely occurred multiple times and independently in different clades of the Viridiplantae. A comparison of the predicted structure of the two *A. thaliana* SmE proteins suggested that two amino acid differences that distinguish PCP and PCPL are unlikely to affect their function. In contrast to mutations in *PCP*, *pcpl* mutant plants did not display any phenotypes under any of the temperatures tested even though *PCPL* gene expression is modulated by temperature. Functional analyses indicate that the two *A. thaliana* SmE proteins can in principle fulfill the same function and differences reside mostly in their transcriptional regulation. We found that the expression of *PCP* is linked to its introns. Among them the 1^st^ and 2^nd^ intron seem to be of particular importance and deletion of either of these introns strongly affected *PCP* expression. However, other introns also seem to contribute to the regulation of *PCP* expression. These findings suggest that *cis*-regulatory elements distributed across multiple introns or complex interaction between non-coding and coding sequences during transcription and RNA processing ensure timely and correct expression of *PCP* in response to temperature.

## Results

### *SmE* duplication is a reoccurring event throughout plant evolution

*A. thaliana* genome encodes two SmE proteins, AtPCP (SmE1; At2g18740) and AtPCPL (AtPCP-like; SmE2; At4g30330). To reconstruct the evolution of the *SmE* paralogs we retrieved *SmE* gene and protein sequences from 24 species of green plants. These represented major lineages such as green algae (*Chlamydomonas reinhardtii*), bryophytes (*Physcomitrium patens*), lycophytes (*Selaginella moellendorffii*,) and ferns (*Ceratopteris richardii*). Most of the SmE genes and proteins, however, belong to the angiosperms, including *Amborella trichopoda* (the sister species to the rest of angiosperms) as well as several monocotyledonous species and representatives of two major Eudicot clades, namely the Asterids and Rosids. Five Brassicaceae species were added to the analysis to increase local resolution in proximity to *A. thaliana*. Out of these 24 species, two, *Chlamydomonas reinhardtii* and *Ceratopteris richardii*, possessed only one *SmE* gene while in the remaining species between 2 to 6 paralogs of *SmE* could be identified (Table S1).

Initial alignment of the genomic DNA sequences showed a very high conservation within the exon sequences only. The low sequence similarity of the introns as well as a big variability of their lengths prevented the construction of a reliable phylogenetic tree based on the genomic DNA alignment. Instead, we proceeded with reconstructing a SmE phylogeny using amino acid and cDNA sequence alignments. The phylogenetic tree overall mapped to the currently accepted evolutionary tree of angiosperms (Zuntini et al. 2024) (Fig. 1). *SmE* genes did not split into two distinct groups, *SmE1-like* (including *AtPCP*) and *SmE2-like* (including *AtPCP-like*), rejecting our initial hypothesis that *A. thaliana PCP* and *PCPL* were the results of an ancient gene duplication in the green lineage. Instead, our results suggest that *SmE* duplications occurred multiple times independently in various branches of the green lineage and that the *SmE* duplication that led to *AtPCP* and *AtPCPL* appears to be specific to the Brassicaceae family. Interestingly, we did not observe *SmE2-like* genes in *Brassica rapa* and *Brassica oleracea*, which instead contained additional copies of *SmE1*, suggesting the secondary loss of *PCPL-like* genes and additional *PCP* duplication in the *Brassica* genus. Similar duplication events that were inherited by a whole clade of closely related species occurred in grasses (*Poaceae*) and presumably in the *Solanaceae* family. Duplications observed in the remaining examined species did not group inside clades above the species level.

**Fig. 1.**
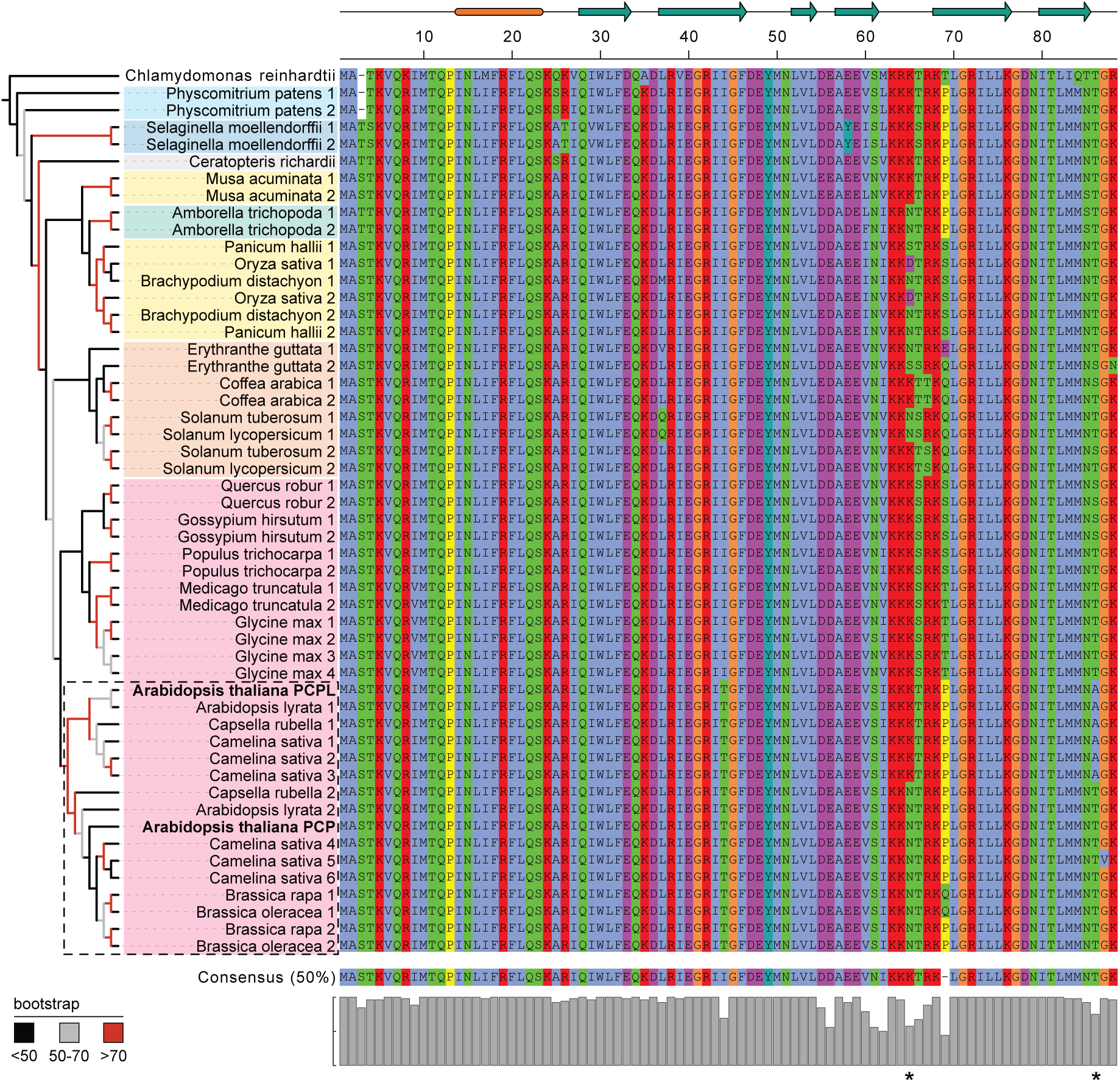
Phylogenetic tree of *SmE* genes and corresponding SmE protein alignment. A maximum likelihood tree based on the CDS alignment was constructed using IQ-TREE software. Nodes are colored according to the bigger clades they belong to: green algae (no color), Bryophytes (light blue), Lycophytes (dark blue), Ferns (grey), Monocots (yellow), Eudicots (orange for Asterids and magenta for Rosids). The Brassicaceae family is highlighted in a dashed box. Protein sequences are colored according to the ClustalX color scheme (Thompson et al. 1997). *SmE* copies are numbered for each species except *Arabidopsis thaliana*, where gene names *PCP* and *PCPL* are given instead and highlighted in bold. Corresponding gene IDs can be found in Supplementary Table S1. The schematic drawing above the protein alignment represents the secondary structure of the protein. The N-terminal alpha-helix is indicated in orange and teal arrows mark beta-strands. Asterisks mark amino acid positions 65 and 86 that discriminate PCP (SmE1) and PCPL (SmE2).

### Brassicaceae-specific amino acid substitutions are unlikely to affect SmE protein function

Our phylogenetic analysis indicated that *A. thaliana PCP* and *PCPL* are the result of a Brassicaceae-specific duplication. Specifically, the SmE1-like and SmE2-like proteins differ in two amino acids, an asparagine to lysine substitution in position 65 and a threonine to alanine substitution in position 86. The question arises if these amino acid substitutions affect the function of the proteins.

Both amino acids map to regions that are unlikely to play crucial roles in protein folding and affect its function. Position 65 falls into the loop region between two beta-strands or the very beginning of the beta-strand, and position 86 lays in the unstructured C-terminal region of the protein after the final predicted beta-strand. Furthermore, substitutions in position 65 are quite diverse in their physical properties and occurred independently several times in the green lineage (Fig. 3). In contrast, substitutions in position 86 are not as widespread, but the substitution of threonine (T) in PCP by alanine (A) in PCPL could in principle affect post-translational protein modifications, in particular phosphorylation. Nevertheless, considering the position and nature of the two amino acid substitutions they seem unlikely to affect protein function. This notion is supported by the predicted structures of PCP and PCPL, which are largely congruent (Fig S1). In particular, the substitutions in positions 65 and 86 do not appear to affect the protein backbone and structure (Fig. S1).

### SmE proteins are essential for plant development

Loss of PCP (SmE1) in *A. thaliana* has been reported to result in temperature-sensitive phenotypes. *pcp-1* was initially described as a mutant sensitive to cold (4°C) and cool ambient temperature of 16°C (Huertas et al. 2019; Capovilla et al. 2018). In agreement with these previous reports, we observed an intermediate phenotype with drastic shortening of the root and anthocyanin accumulation in the cotyledons in *pcp-1* grown at 10°C (Fig. S3). We also observed a mild delay in the development of 10-days-old *pcp-1* at 23°C. Mutants had smaller leaf rosettes and shorter roots with less abundant and developed lateral roots in comparison to Col-0 (Fig. S4). This delay in development appeared to be transient, and *pcp-1* plants eventually caught up and developed similarly to WT at 23°C (Capovilla et al. 2018). Confirming its cold temperature-sensitive nature, 7-day-old *pcp-1* seedlings grown at 27°C did not exhibit significant alterations in shoot and root development compared to Col-0 plants growing in the same conditions (Fig. S3).

Consistent with the cold-sensitivity of the *pcp-1* mutants, *PCP* expression in the WT was induced two-fold at 16°C and almost 3.5-fold at 10°C in comparison to 23°C (Fig. 2 A). We also observed a mild induction of *PCP* in response to elevated temperature. Interestingly, our qPCR data showed that *PCPL* in WT displayed the same overall expression profile as *PCP* (Fig. 2 B) but the expression changes in response to cool and warm temperatures were much weaker. For example, whereas *PCP* expression was induced more than 3-fold at 10°C when compared to 23°C, *PCPL* expression increased only 1.5-fold. These findings suggested that *PCP* might be the more important of the two *SmE* genes in response to temperature.

**Fig. 2.**
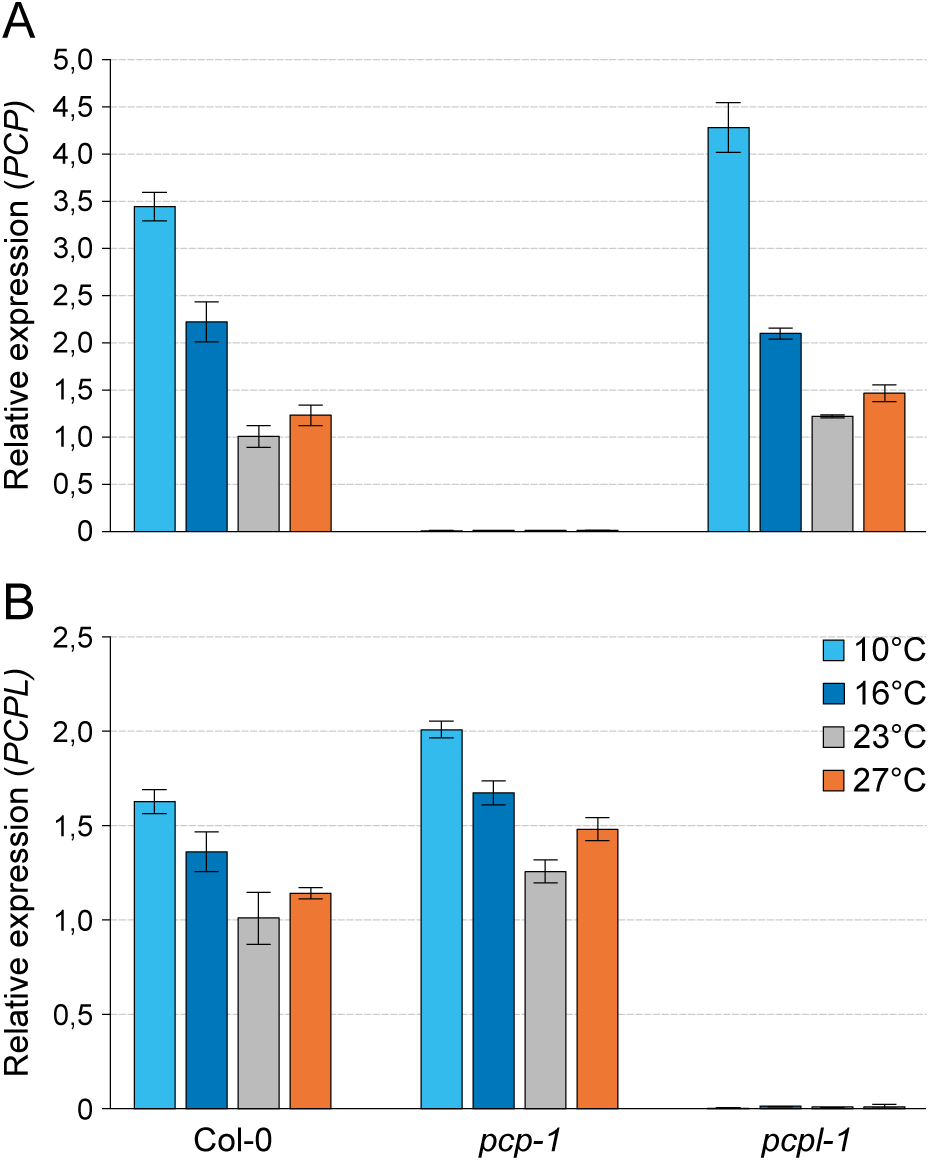
Expression of *PCP* and *PCPL* in *A. thaliana* SmE mutants. *PCP* (SmE1) (A) and *PCPL* (SmE2) (B) expression was determined in the Col-0, *pcp-1*, and *pcpl-1* grown at 10°C, 16°C, 23°C and 27°C by RT-qPCR. Bars show the mean expression based on 3 biological replicates with 3 technical replicates each. Error bars indicate the standard deviation (SD).

To test this hypothesis, we generated two *pcpl* mutant alleles using CRISPR/Cas9 mutagenesis. Interestingly, we found that *pcpl-1* and *pcpl-2* mutants were indistinguishable from wild-type across a wide range of temperatures (Figs. S3, S4), which is remarkably different from the temperature-sensitive *pcp-1* (Fig. S3, S4). Possibly, the expression of *PCPL* is sufficient to compensate for the loss of *PCP* in *pcp-1* at 23°C and higher temperatures, but *PCPL* is not enough to fulfill the splicing needs of the cell at lower temperatures. Alternatively, there might be differences in the processing of the transcripts or even in the proteins themselves.

However, plants clearly require at least one functional copy of *SmE* as demonstrated by the finding that 25% of the developing seeds of self-pollinated double-heterozygous *pcp pcpl* plants were aborted (Fig. 3), indicating gametophytic lethality. The same experiment also revealed a moderately increased seed abortion rate in the *pcp-1* single mutant. In contrast, the seed set was essentially unaffected in the *pcpl* single mutants (Fig. 3), again highlighting phenotypic differences between the mutants in these paralogous genes.

**Fig. 3.**
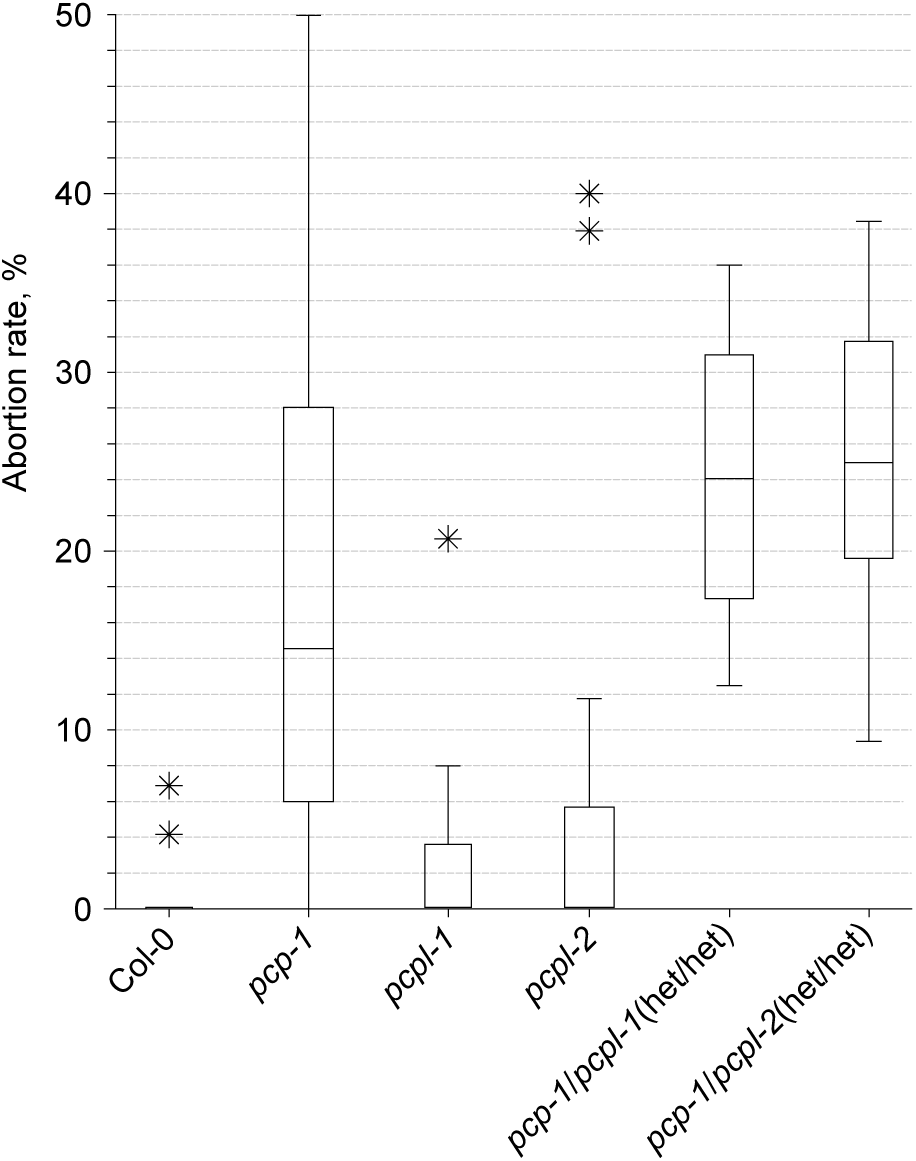
Abortion rate of the SmE mutants. The proportion of the aborted ovules to the total amount of ovules in the silique was calculated (n = 20) and plotted as a box-with-whiskers plot with Tukey test. The box spans over the 25^th^ to 75^th^ percentiles, the line inside the box represents the median, whiskers reach out to the minimal and maximal values in each dataset. Asterisks marks the outliers.

### PCP expression requires genomic and intronic sequences

Since our phylogenetic and structure-function analyses did not reveal any major differences in PCP and PCPL proteins, the question arises as to why the two mutants have such different phenotypes. Hypothesizing that the reason might reside in the transcriptional regulation of the genes, we created constructs expressing either the CDS or the genomic DNA (gDNA, i.e. exons plus introns) of *PCP* and *PCPL* under the regulation of the *PCP* promoter and terminator sequences. In addition, we also generated constructs overexpressing the CDS of either of these two genes under the control of the constitutive *p35S* promoter (Table S2).

We introduced these constructs into the *pcp-1* mutant and used primary root length and overall morphology as a measure of phenotypic rescue. Interestingly, at temperatures between 10°C to 23°C lines (#1-3) carrying the *pPCP::cPCP::tPCP* construct exhibited a *pcp-1*-like phenotype, which got more severe at lower temperatures (Fig. 4 A, S3, S4). At the same time, *p35S*-mediated overexpression of the *PCP* CDS resulted in full phenotype restoration (Fig. 4 B, S3, S4). Similarly, *pPCP::gPCP::tPCP* constructs (#7-9) fully rescued the *pcp-1* phenotype (Fig. 5, S3, S4). Consistent with the phenotyping data, RT-qPCR showed that in the case of the gDNA and overexpression lines *PCP* expression was restored to approximately 70% of the WT levels and strongly overexpressed, respectively (Fig. 6 A). Importantly, the expression of both *pPCP::gPCP::tPCP* and *35S::cPCP::tPCP* also restored the temperature-dependent expression of *PCP*, which might indicate post-transcriptional regulation in addition to transcriptional regulation via *cis*-regulatory elements present in the genomic sequence of PCP. In contrast, no expression of *PCP* beyond the residual expression typical for the *pcp-1* mutant could be detected in the lines expressing the *PCP* CDS under the control of *pPCP* and *tPCP*. Taken together, these findings support the idea that phenotype restoration correlates with the expression of *PCP*, and that the full gDNA sequence is essential for the proper expression.

**Fig. 4.**
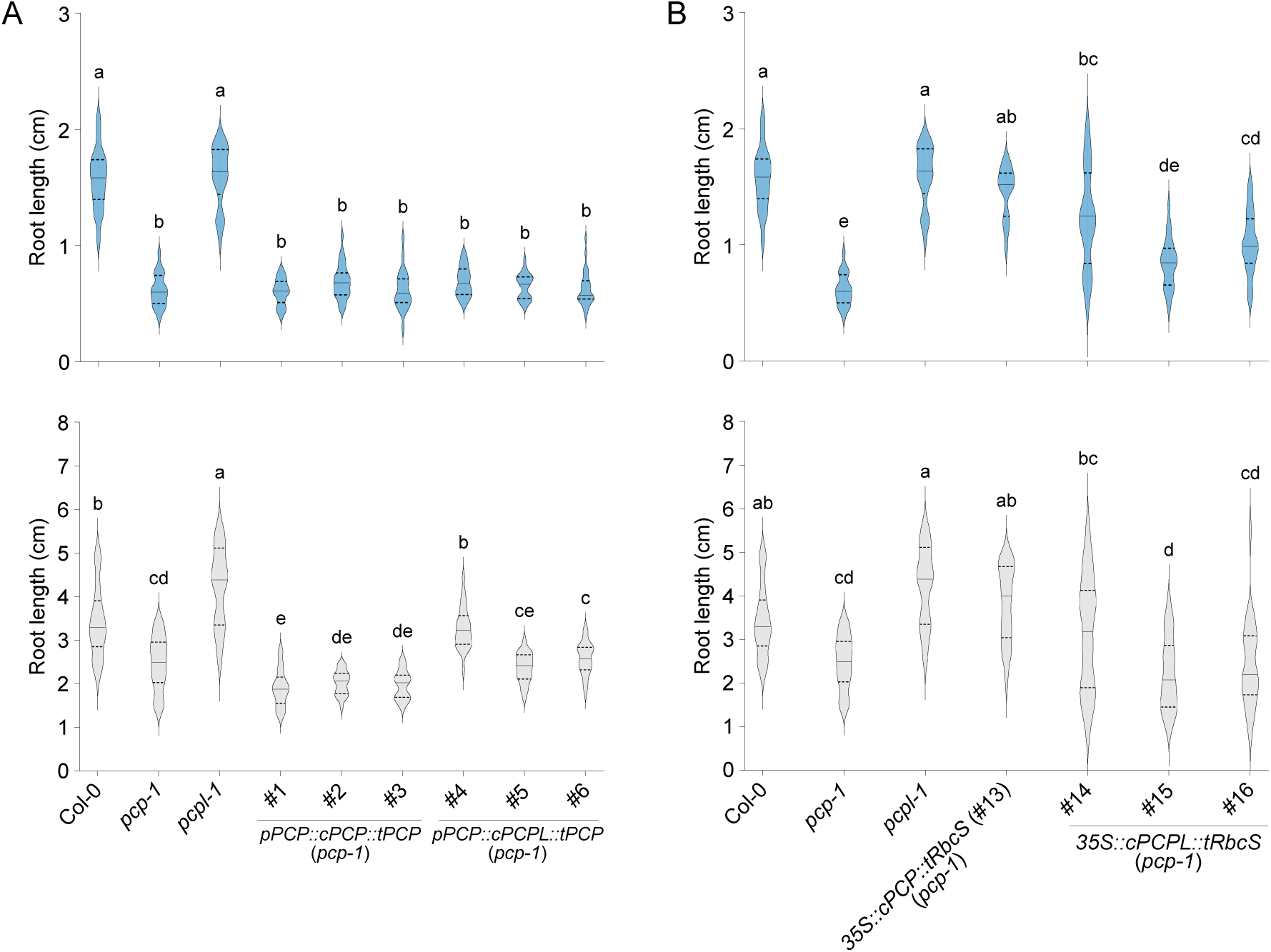
Root length of lines expressing SmE coding sequences. (A) Root phenotype of three independent lines carrying *pPCP::cPCP::tPCP* (#1-3) and *pPCP::cPCPL::tPCP* (#4-6) constructs in the *pcp-1* background. Col-0, *pcp-1*, and *pcpl-1* serve as controls. (B) Root phenotype of three independent lines carrying *35S::cPCPL::tRbcS* (#14-16) constructs in the *pcp-1* background. Col-0, *pcp-1*, *pcpl-1*, and previously established *35S::cPCP::tRbcS* (#13) (Capovilla et al. 2018). The length of the primary root of 10-day-old seedlings grown at 16°C (blue) and 23°C (light grey) is shown as violin plots (n = 20 to 25). Solid lines represent the median and dashed lines represent quartiles. One-way ANOVA test with Tukey correction was performed using GraphPad Prism. Letters represent significantly different (P < 0.05) groups.

**Fig. 5.**
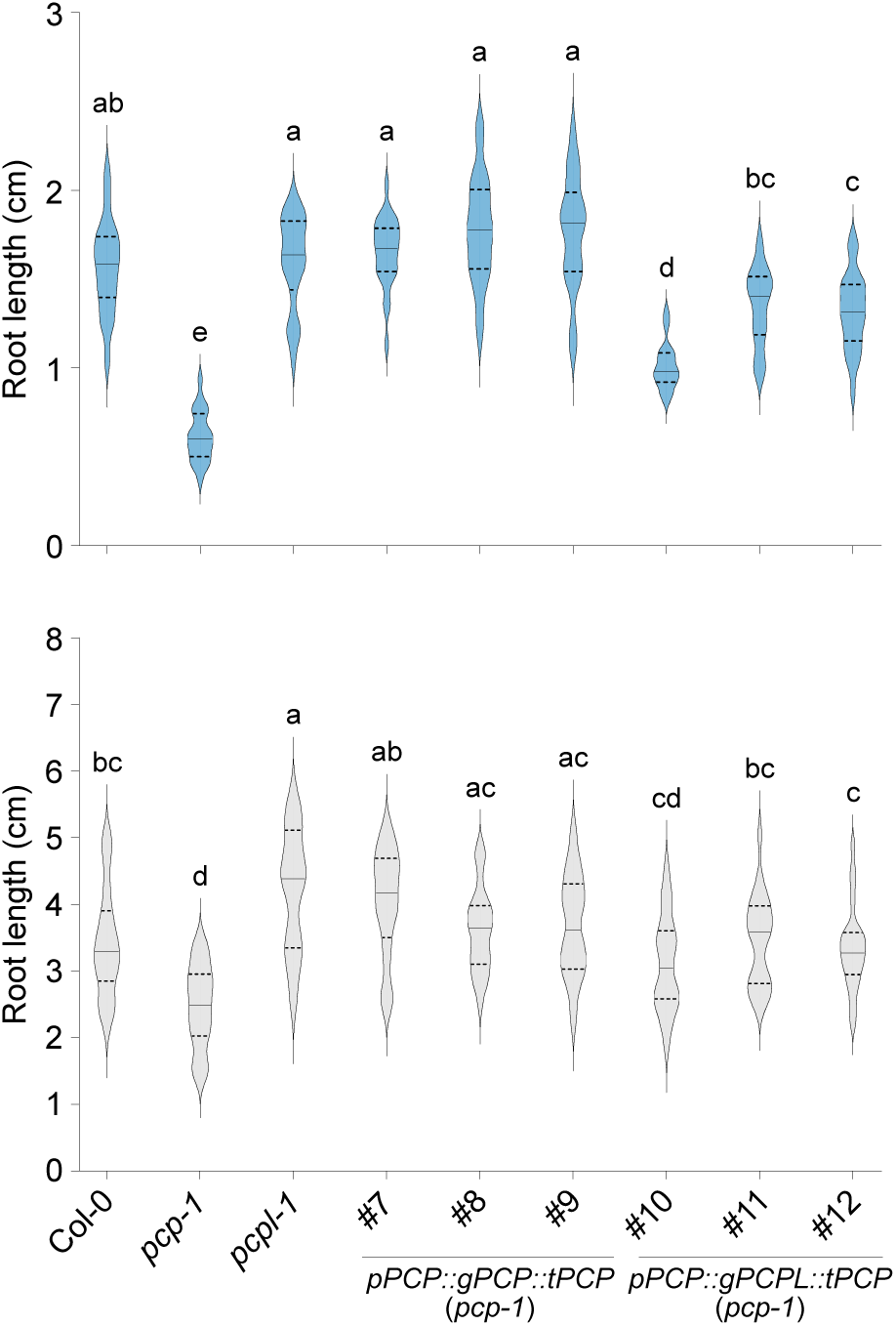
Genomic *PCP* construct rescues the root length phenotype of *pcp-1*. Length of the primary root of three independent lines in the *pcp-1* background transformed with *pPCP::gPCP::tPCP* (#7-9) and *pPCP::gPCPL::tPCP* (#10-12) constructs. Col-0, *pcp-1*, and *pcpl-1* serve as controls. Root length was determined using 10-day-old seedlings grown at 16°C (blue) and 23°C (light grey). Root lengths are shown as violin plots (n = 20 to 25) where a solid line represents the median and dashed lines represent the quartiles. A one-way ANOVA test with Tukey correction was performed using GraphPad Prism. Letters represent significantly different (P < 0.05) groups.

**Fig. 6.**
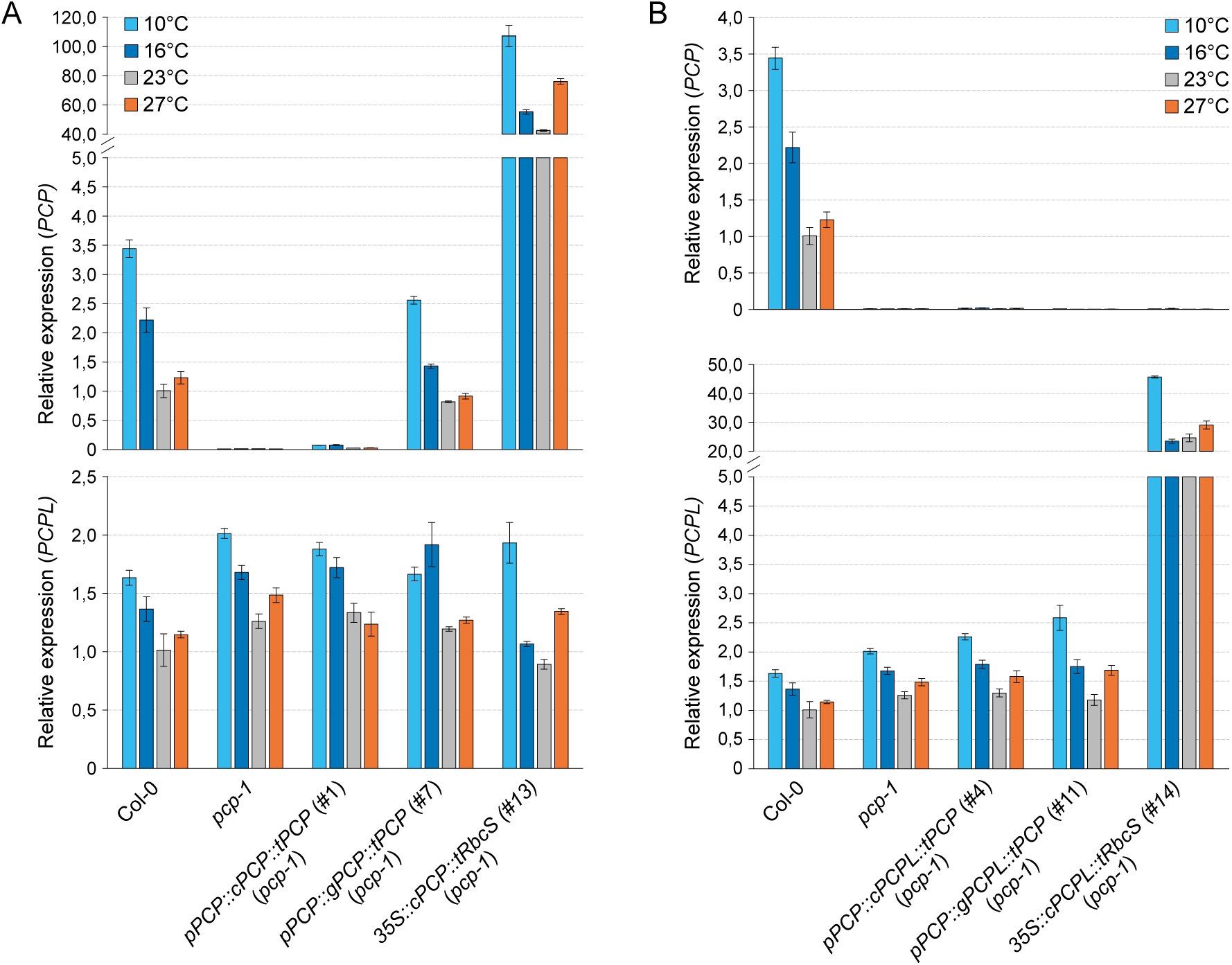
Expression analysis of *PCP* and *PCPL*. (A) Expression of *PCP* and *PCPL* in Col-0, *pcp-1*, and lines carrying *pPCP::cPCP::tPCP* (#1), *pPCP::gPCP::tPCP* (#7), and *35S::cPCP::tRbcS* (#13) (Capovilla et al. 2018a) constructs in the *pcp-1* background. (B) Expression of the *PCP* and *PCPL* in Col-0, *pcp-1*, and *pcp-1* lines carrying *pPCP::cPCPL::tPCP* (#4), *pPCP::gPCPL::tPCP* (#11), and *35S::cPCPL::tRbcS* (#14). Gene expression was determined in seedlings after 5 days of growth at 23°C followed by 24 h of growth at 10°C (light blue), 16°C (dark blue), 23°C (grey), and 27°C (orange). Bars show the mean expression calculated from 3 biological replicates with 3 technical replicates each. Error bars indicate the standard deviation (SD).

### Transcription regulation elements differ between PCPL *and* PCP

Similarly, expression of the *PCPL* CDS under the control of the *pPCP* and *tPCP* (lines #4-6) was unable to rescue the *pcp-1* phenotype at 16°C and 10°C (Fig 4 A, S3, S4). Except for line #4, which showed a moderate phenotype rescue, all lines were statistically indistinguishable from *pcp-1* also at 23°C, while RT-qPCR analysis detected mild elevation in the *PCPL* expression at all temperature conditions (Fig. 6 B).

Lines #10-12, which expressed the *gPCPL* construct including all introns under the control of the *PCP* promoter and terminator (*pPCP::gPCPL::tPCP*) showed partial to full phenotypic rescue of the *pcp-1* phenotype at 16°C and 23°C (Fig. 5, S3). However, irrespective of temperature (10°C to 23°C) these lines performed worse than their counterparts (lines #7-9) expressing *gPCP* under the control of the *PCP* promoter and terminator (Fig. 5 S3, S4). Expression of *PCPL* in line #11 was upregulated in comparison to Col-0 at all temperatures, especially at 10°C. *PCPL* expression did not differ significantly between #11 and *pcp-1* except at 10°C (Fig. 6 B). Together, these findings suggest a discrepancy between the regulation of *PCP* and *PCPL* expression, likely due to different regulatory elements in the gDNA sequences of *PCP* and *PCPL*.

Overexpression of *cPCPL* (lines #14-16) in the *pcp-1* background resulted in partial to full restoration of the root morphology. However, overall phenotype of the rescue lines was more variable than in the WT plants and in the other rescue lines (Fig. 4 B). Possibly, this variability is a result of the very high overexpression of *PCPL* in these lines, which vastly exceeds the endogenous *PCPL* levels. Such misregulation of a central splicing factor can be expected to have unwanted effects on downstream genes, which could result in the observed high phenotypic variability.

### PCP and PCPL proteins are functionally redundant

Having established a genomic *PCP* construct that not only rescued the phenotype of *pcp-1* at low temperatures but also the temperature-responsive expression enabled us to address the role of the amino acid substitutions that discriminate PCP and PCPL proteins. To this end we introduced the amino acid substitutions found in PCPL (N65K; T86A) into the *gPCP* genomic rescue construct (*pPCP::gPCP_N65K_T86A::tPCP*; lines #19-20) and introduced it into *pcp-1*. We found that this construct fully restored the WT phenotype at 16°C (Fig. 7) and 23°C, confirming our previous assumptions that the SmE1 and SmE2 proteins in *A. thaliana* were functionally redundant.

**Fig. 7.**
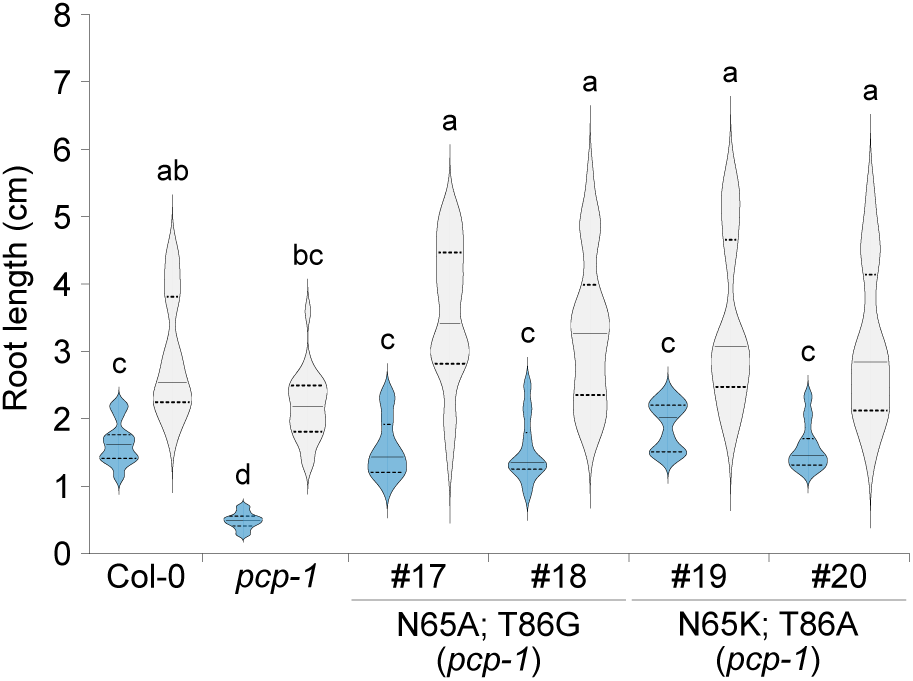
Root length of PCP-PCPL amino acid substitution lines. Length of the primary root of plants carrying *pPCP::gPCP_N65A_T86G::tPCP* (#17-18) and *pPCP::gPCP_N65K_T86A::tPCP* (#19-20) rescue constructs in the *pcp-1* background. Col-0 and *pcp-*1 served as controls. Root length was measured using 10-day-old seedlings grown at 16°C (blue) and 23°C (light grey). Root lengths are shown as violin plots (n = 20 to 25) where the solid line indicates the median and the dashed lines represent the quartiles. A one-way ANOVA test with Tukey correction was performed using GraphPad Prism. Letters represent significantly different (P < 0.05) groups.

We also prepared a similar construct resulting in the SmE protein with alanine and glycine substitutions (*pPCP::gPCP_N65A_T86G::tPCP; lines* #17-18), to test the importance of the amino acids in positions 65 and 86 for SmE function. Expressed in the *pcp-1* background this construct also fully restored its phenotype at both temperatures tested, suggesting that the amino acids in positions 65 and 86 play only a minor role in SmE function.

### Introns work together to regulate PCP expression

Since the full genomic sequence of *PCP* with 5’ promoter and 3’ UTRs was necessary for complete *pcp-1* phenotype rescue, and expression of the *PCP* cDNA under the *pPCP* and *tPCP* did not result in expression of the gene, we assumed that *PCP* expression was linked to its introns. To test this hypothesis, we created several *gPCP* intron-deletion constructs (Table S2, S3), introduced them in *pcp-1*, and determined the extent of phenotypic rescue (Fig. 8).

**Fig. 8.**
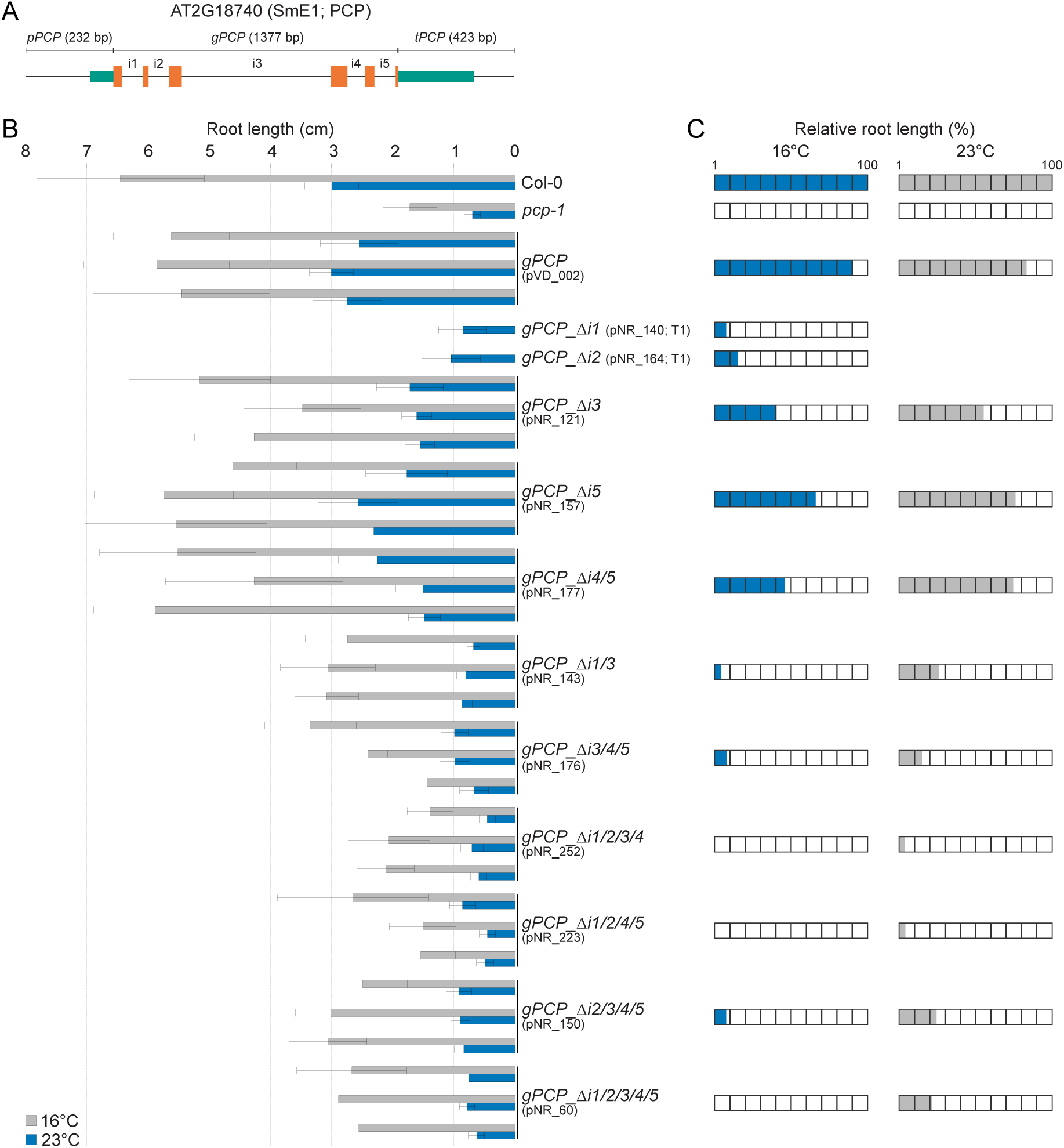
Introns are required for *PCP* expression. (A) Schematic representation of the full genomic *PCP* construct used to perform the intron deletion studies. Teal boxes represent promoter and terminator regions, orange boxes represent exons and solid lines represent non-coding sequences. i1 to i5 labels indicate introns 1 to 5. (B) Length of the primary root of three independent lines in the *pcp-1* background transformed with various intron deletion constructs (left). Deletion lines and constructs are labeled as *gPCP_ΔiX*, where Δ stands for deletion, i stands for intron, and X indicates the number(s) of the deleted intron(s). Col-0, *pcp-1*, and a *pcp-1* rescue line carrying the full-length *pPCP::gPCP::tPCP* construct serve as controls. Root length was determined using 10-day-old seedlings grown at 16°C (blue) and 23°C (light grey). Bars show the mean of the root length measurement (n = 11 to 37), and error bars indicate the standard deviation (SD). (C) Average root-phenotype rescue relative to Col-0 (100%) and *pcp-1* (0%) based on the root length of the three independent lines shown in (B).

Interestingly, the deletion of the largest intron, intron 3 (*gPCP-Δi3*), which we had speculated might contain essential *cis*-regulatory elements, resulted in a partial rescue of the *pcp-1* root phenotype (Fig. 8 B). Similarly, the deletion of intron 5 (*gPCP-Δi5*) alone as well as the combined deletion of intron 4 and 5 (*gPCP-Δi4/5*), resulted in partial phenotype rescue. In both cases phenotype rescue was slightly better than that of (*gPCP-Δi3*). These findings suggest that even though these introns likely contribute to *PCP* function, they are by themselves not essential for *PCP* regulation. However, deletion of all three introns together (*gPCP-Δi3/4/5*) had an additive effect and resulted in a *pcp-1*-like phenotype. Deleting an additional other intron had a similar effect. In contrast, deletion of either intron 1 (*gPCP-Δi1*) or intron 2, (*gPCP-Δi2*), almost completely abolished phenotypic rescue of *pcp-1* at 16°C, indicating that these introns are essential for *PCP* expression. Similarly, any higher-order intron deletion construct missing either intron 1, intron 2, or both failed to rescue the *pcp-1* mutant. On the other hand, the presence of either intron 1 or intron 2 in the CDS alone was insufficient to drive the expression and restored the *pcp-1* phenotype only partially. Taken together, our results highlight the importance of introns 1 and 2 for *PCP* activity but also indicate that introns 3-5 contribute as construct lacking these three introns are unable to rescue *pcp-1*. Whether introns 1 and 2 contain specific *cis*-regulatory elements essential for *PCP* expression or if they boost expression by other means such as promoting transcription in general, or processing of the pre-mRNA remains to be established.

## Discussion

Throughout the course of evolution, plants underwent multiple genome duplications and preserved copies of paralogous genes. Through neo- and sub-functionalization these paralogues either acquired new functions or performed their original tasks under different conditions, thereby increasing the adaptability of plants to adverse environmental conditions (Flagel and Wendel 2009).

Here we investigated *SmE* genes and proteins to shed light on the evolution of this important regulator of RNA splicing in the green lineage. Consistent with the work by Cao et al. 2011 who had analyzed the relationship among Sm proteins in *A. thaliana*, we found that the duplication resulting in *PCP/PCPL* happened independently of *SmE* duplications in other plant clades during the most recent Brassicaceae-specific whole-genome duplication (WGD) event, namely At-α (Barker et al. 2009). In general, reoccurring duplications of *SmE* could be beneficial for plant adaptation, as additional copies could ensure sufficient expression under specific conditions, which might explain a wide range of biotic and abiotic stresses that affect *SmE* expression across different species of the Viridiplantae. Considering the importance of SmE for plant survival, duplications also ensure that at least one functional copy of the gene is always active, while others are available for subfunctionalization (Blanc and Wolfe, 2004).

Despite multiple independent duplication events in various clades of the Viridiplantae, the *SmE* CDS and the protein sequences exhibit a very high degree of sequence conservation. This is in line with a study by Chen and Cao 2014, who reported a significant prevalence of synonymous over nonsynonymous mutations during *SmE* evolution. Interestingly, the same study also showed that the exon-intron structure of homologous *Sm* genes was conserved even between distantly related species. In agreement with this report, we observed the same 6 exon / 5 intron structure found in *AtPCP* in euphyllophytes (ferns and seed plants). In contrast, the representatives of green algae, bryophytes and lycophytes were characterized by single-exon *SmE* coding sequences.

Cao et al. 2011 also reported that duplicated *Sm* genes showed little similarity in their expression patterns despite the high conservation in their intron-exon structure. However, in the case of *AtPCP* and *AtPCPL*, analysis of publicly available expression data indicates their co-expression across a wide range of tissues and treatments. In line with this finding, we observed that the expression of *PCP* and *PCPL* in response to ambient temperatures ranging from 10°C to 27°C is quite similar. Despite these similarities in temperature-dependent gene regulation, we did not observe any temperature-sensitive phenotype in *pcpl* mutants (Fig. 4, S3, S4).

Interestingly, rice *SmE* orthologs were shown to be induced in response to cold (Chen and Cao 2014;Kawahara et al. 2016) reminiscent of *PCP* in *A. thaliana*. Similarly, it has been reported that *SmE* genes in tomato and *Populus* were also differentially expressed in response to cold (Chen et al. 2015;Barrero-Gil et al. 2016;Filichkin et al. 2018). In contrast, *SmE* homolog in *C. reinhardtii,* which is encoded by a single-exon gene, was not induced in response to cold (Li et al. 2020b). This finding could be interpreted as evidence that expression regulation by cold temperatures is a derived feature and might be conserved in higher land plants. However, considering that gene duplications provide a source for inter- and intraspecies expression and functional diversification (Gu et al. 2004; Ha et al. 2009) additional analyses, including the Brassicaceae *SmE* homologs, are required before any conclusions regarding the contribution of the *SmE* gene structure expression in response to cold can be drawn.

Further investigation of the protein structure of the *A. thaliana* SmE proteins suggested that the two proteins did not functionally diversify. This notion is supported by our finding that substituting the two amino acids that discriminate PCP and PCPL in the context of the full genomic *PCP* rescue construct by either the amino acids found in PCPL or two other small amino acids resulted in the full rescue of the *pcp-1* phenotype at 16 °C (Fig. 7). This is in agreement with the observation that gene copies retained after the WGD show lower levels of structural and functional changes in the course of evolution than tandem duplications (Wang et al. 2013). Taken together, these findings suggest that the main difference between *PCP* and *PCPL* resides in the differential regulation of the expression of these two genes. In *A. thaliana*, *PCP* might be the more important orthologue, especially at low temperatures, not only due to the differential expression in response to cold but also due to the downstream regulation of the transcripts (Fig. 6). As mentioned above shift in expression patterns between duplicated genes contribute to the retention of the duplicates (Duarte et al. 2006; Blanc and Wolfe 2004). Furthermore, the fact that *Brassica rapa* and *Brassica oleracea* have lost their copy of *PCPL* and acquired an additional copy of *PCP* can be interpreted as indirect evidence for the functional redundancy of the two proteins, with PCP being the more important in *A. thaliana.* However, abortion rate of 25% in the *pcp-1 pcpl* crossing progeny suggests that plants require at least one functional copy of SmE for survival (Fig. 3). It has previously been demonstrated that PCP is important for proper splicing of pre-mRNA (Huertas et al. 2019; Hong et al. 2021; Wang et al. 2022; Hong et al. 2023) but information regarding target preferences of PCPL is still lacking, and we cannot completely exclude the possibility that the two *A. thaliana* SmE proteins interact with different partners or are differentially regulated at some level.

In this context, it is important to note that our results indicate that expression of *PCP* requires a full genomic sequence including promoter, 5’ UTR, gene region consisting of exons and introns, and downstream sequences. Only such a full genomic construct can rescue the *pcp-1* phenotype at low ambient temperature and temperature-responsive expression, while expression of the *PCP* cDNA under the control of the *PCP* promoter and terminator cannot (Fig. 5, 4 A). Expression of the *PCPL* gDNA under the control of the *PCP* promoter and terminator, despite being highly similar to *PCP* in exon-intron structure, was not sufficient to completely rescue the *pcp-1* mutant (Fig. 5). It should be noted, that Wang et al. 2022 found that expression of a *pPCP::gPCP-3xFLAG* construct lacking the terminator was able to rescue the *pcp-1* phenotype, which suggests that minimal sequence requirements for *PCP* expression do not include the 3’ region. Our intron deletion experiments clearly demonstrate that intronic sequences are absolutely required for *PCP* expression and function (Fig. 8). In particular, the first two introns seem to play major roles in promoting *PCP* expression as deletion of either of them renders the *PCP* rescue construct non-functional. However, the 1^st^ intron on its own (*gPCP-Δi2/3/4/5*) is not sufficient to drive the expression of *PCP* to rescue the *pcp-1* mutant phenotype (Fig. 8). Furthermore, successive deletion of other introns 3, 4, and 5 had a negative effect on *PCP* expression, highlighting the important role introns play in the regulation of *PCP* expression.

Introns show a very wide range of regulatory mechanisms that affect not only gene expression but also the processing of the transcript, its metabolism, and even translational efficiency, which complicates the study of intron-mediated expression regulation (Le Hir et al. 2003; Chorev and Carmel 2012). Introns can drive the expression of the gene, affect expression levels and organ-/tissue-specificity, activate transcription, contain *cis*-regulatory elements, provide miRNA-binding sites, and possibly contain elements that act as riboswitches (Xie and Wu 2002; Meng et al. 2013; Meng et al. 2021). First introns in particular have been shown to play important roles in these regulatory functions, in some cases turning them into the key regulators of gene expression (Salgueiro et al. 2000; Zalabák and Ikeda 2020; Zhu et al. 2024). In particular, the phenomenon of the intron-mediated enhancement (IME) has been shown to be strongly linked to the proximity of the given (first) intron to the 5’UTR (Rose 2004; Gallegos and Rose 2015).

Irrespective of the mode by which introns promote the expression of *PCP*, other factors have been shown to contribute to regulating *PCP*. For example, induction of *PCP* in response to cold was found to be attenuated in a *cbf123* triple mutant background after 12 h or more (Z. Wang et al. 2022), indicating that these key regulators of plant temperature responses contribute to *PCP* regulation. Notably, this regulation appears to be direct as EMSA indicates interaction between the promoter region of *PCP* and CBFs and the presence of a CRT/DRE *cis*-regulatory element (Wang et al. 2022). However, regulation of *PCP* by the CBFs is likely not a rapid response as indicated by the fact that its expression was unchanged 6 h after exposure to 4 °C (Huertas et al. 2019).

In summary, our work provides evidence that SmE is a protein essential for plant growth and development and that the two copies of the SmE protein present in *A. thaliana* perform the same function. However, in contrast to *PCP*, the deletion of *PCPL* does not affect plant growth and development under any conditions tested. We attribute these differences to divergence in gene expression as *PCP* is in general the more highly expressed of the two genes. Interestingly, our results indicate that intronic sequences are central to *PCP* expression. The molecular mechanism underlying this regulation is, however, currently unclear and will need to be investigated further in the future.

## Materials and methods

### Phylogenetic analysis

At2g18740 (AtPCP/AtSmE1) was used to query the Phytozome v13 (Goodstein et al. 2012) database to identify potential homologues among the available genomes of Viridiplantae. Species were chosen to represent the major groups of the green plants as best as possible (Supplementary Table S1). Genomic DNA, coding DNA, and protein sequences of selected SmE orthologs were downloaded from the NCBI GenBank database (www.ncbi.nlm.nih.gov/genbank/).

The retrieved SmE amino acid sequences of the selected species were aligned using MAFFT v7 (L-INS-i algorithm) (Katoh and Standley 2013) and manually checked for mistakes in the alignment using Geneious Prime 2023.0.4 software (https://www.geneious.com/). The corrected amino acid alignment was reverse translated into a corresponding cDNA alignment using Pal2Nal v14 (Suyama et al. 2006). The output cDNA alignment was used to build a phylogenetic tree using IQ-TREE v2.2.0 (Minh et al. 2020; Kalyaanamoorthy et al. 2017). The tree was rooted on the green alga *Chlamydomonas reinhardtii.* Command lines used to perform the tree calculation are provided in Supplementary Method S1. Trees were visualized using the iTOL online tool v6 (Letunic and Bork 2007).

### Protein structure prediction

Protein domain search and secondary structure prediction were performed on the consensus sequence reconstructed after protein alignment using NCBI Conserved Domains Search (Marchler-Bauer et al. 2017; Lu et al. 2020; Wang et al. 2023) and Jpred 4 tool (Drozdetskiy et al. 2015), respectively. 3D protein structures of PCP (SmE1) and PCPL (SmE2) based on AlphaFold2 predictions (Jumper et al. 2021) were retrieved from the BAR ePlant (https://bar.utoronto.ca/eplant/) (Waese et al. 2017). Protein structures were visualized using PyMOL, version 3.0.2 (Schrödinger, LLC).

### Plant material and growth conditions

All work was done using *Arabidopsis thaliana* plants of Col-0 accession. The *pcp-1* T-DNA insertion line (SALK_089521) has previously been described (Capovilla et al. 2018). The *pcpl-1* and *pcpl-2* deletion mutants were generated as part of this study. Briefly, *A. thaliana* Col-0 plants were transformed by floral dipping with constructs for CRISPR/Cas9-mediated mutagenesis (described below) (Clough and Bent 1998). Primary transformants (T1) carrying deletions in *PCPL* were identified, and homozygous transgene-free *pcpl-1* mutants were isolated in subsequent T2/T3 generations.

Seeds were stratified at 4°C in the dark for 48 h before sowing on soil or plates. Seeds grown in sterile conditions were sown on ½ MS media (Murashige and Skoog 1962) plates with 1.6% agar. Plates for the selection of transformants were additionally supplemented with 10 ug/mL phosphinotricin (Duchefa Biochemie) (BASTA). Seeds sown on plates were surface sterilized beforehand with 0,015% Triton X-100 solution in 70% EtOH for 10 minutes, washed once with 96% EtOH and three times with MQ water.

Plants were grown in long day (LD) conditions (16 h light/8h dark) in Percival chambers equipped with full range LED illumination with the following conditions: light intensity of 120 μmol*m^-2^s^-1^, constant humidity (RH 65%) at either 10°C, 16°C, 23°C, or 27°C.

### Plant transformation and selection of transformants

Electro-competent *Agrobacterium tumefaciens* GV3101 cells containing the pSOUP plasmid required for the replication of pGreen-IIS-based vectors (Hellens et al. 2000; Lampropoulos et al. 2013) were transformed with the plasmids described below and grown on selective media using the appropriate antibiotics. *A. thaliana pcp-1* mutants were grown at 23°C till flowering stage before siliques started to develop. Plants were transformed by floral dipping using *Agrobacterium tumefaciens*-mediated gene transfer (Clough and Bent 1998). T1 seeds were selected by growing on ½ MS media plates with 1.6% agar supplemented with 10 ug/mL BASTA. Seedlings exhibiting BASTA-resistant phenotype were transferred to soil and the presence of insertion or deletion was confirmed by PCR (Supplementary Table S4). Alternatively, the selection of transformants expressing the mCherry or GFP seed marker was performed by picking T1 fluorescent-positive seeds using an epifluorescent stereomicroscope. To remove the CRISPR/Cas9 cassette, plants were backcrossed to Col-0 and transgene-free plants with deletions in *PCPL* were established (Supplementary Fig. S1).

### Plant phenotyping

Phenotypes are reported for homozygous mutants and transgenic lines for which segregation analysis indicated the presence of a single T-DNA insertion unless otherwise stated. Phenotypic analyses were performed using plants grown on ½ MS plates at 10°C, 16°C, 23°C, and 27°C for 25, 10, 10, and 7 days, respectively. Rescue assays of *pcp-1* plants transformed with *PCP* intron deletions were performed in seedlings grown on ½ MS plates at 16°C for 10 days. Plates for root growth assays were scanned using an EPSON Expression 12000XL scanner equipped with SilverFast^®^ 8 software at 1800 dpi resolution or photographed. In addition, pictures were taken using a digital camera to document rosette phenotypes.

Embryo lethality in self-pollinated *pcp-1*/*pcpl-1* (het/het), *pcp-1*/*pcpl-2* (het/het) lines was calculated as the proportion of aborted embryos relative to the total amount of healthy seeds and aborted embryos on one side of the silique.

### Cloning of DNA fragments and assembly of plasmids

Genomic *PCP* and *PCPL* fragments were amplified by PCR using primers listed in Supplementary Table S4 from DNA extracted from Col-0 using the protocol described by (Berendzen et al. 2005) with the following modifications: the volume of the extraction buffer added to the sample before the grinding step was reduced to 500 μl and the incubation time at 95°C was reduced to 1 minute. Intron deletion constructs in *PCP* were created by overlap extension PCR (Horton et al. 1989) using the primers listed in Supplementary Table S4. *PCP* and *PCPL* open reading frames (ORFs) were amplified from cDNA. For this, total RNA was isolated from Col-0 seedlings using the RNeasy^®^ Plant Mini Kit (Qiagen) following the manufacturer’s protocol. Purified RNA was treated with DNase I (Thermo Scientific) followed by first-strand cDNA synthesis using the RevertAid RT kit (Thermo Scientific) with oligo(dT)_18_ primer. The primers used for amplification of *PCP* and *PCPL* Coding Sequences (CDSs) by PCR are listed in Supplementary Table S4.

*PCP* and *PCPL* genomic sequences and ORFs were cloned into GreenGate C-module entry plasmids as previously described (Lampropoulos et al. 2013). *PCP* promoter (232 bp upstream of the start codon, covering the 5’-UTR and intergenic region) and *PCP* terminator (423 bp downstream the stop codon, including the 3’-UTR and intergenic region) sequences were amplified from Col-0 DNA using primers listed in Supplementary Table S4 and cloned into GreenGate entry vectors A and E, respectively. For the generation of *PCP* and *PCPL* overexpression lines, existing A- and E-modules carrying 35S promoter and pea RbcS terminator sequences, respectively, were used (Lampropoulos et al. 2013).

The plant binary plasmids were assembled in destination vector pGGZ003 using GreenGate cloning as previously described (Lampropoulos et al. 2013). Empty B- and D-modules were used in the assembly reaction. BASTA resistance was used as a selection marker in module F in all constructs except *PCP* intron deletion lines, in which case a seed-specific mCherry selection marker was used (Gao et al. 2016; Capovilla et al. 2017; Wu et al. 2018).

Point mutations to exchange amino acids in positions 65 and 86 in PCP and PCPL were generated by PCR using assembled plasmids with *gPCP* (pVD_007) and *gPCPL* (pVD_008) insertions as a template (Supplementary Table S5) and primers listed in Supplementary Table S4.

The final plasmids assembled by GreenGate cloning are listed in Supplementary Table S2. The remaining plasmids were generated by amplifying the inserts using overlap fusion PCR to simultaneously introduce the desired mutations/deletions and A- and G-compatible GreenGate sites, followed by GreenGate cloning into the pGGZ003 destination (see Supplementary Table S3).

For CRISPR/Cas9-mutagenesis of *PCPL*, pairs of gene-specific sgRNAs were designed using CRISPR-P 2.0 (http://crispr.hzau.edu.cn/CRISPR2/) to simultaneously target two positions in the gene (Supplementary Fig. S1). Selected pairs of sgRNAs were introduced into forward and reverse primers (Supplementary Fig. S1 and Supplementary Table S4), which were used to amplify a dual-guideRNA cassette via the pCBC-DT1T2 vector (Xing et al. 2014) thereby introducing a sgRNA scaffold to the first guide RNA sequence, a U6-29 Promoter to the second guide RNA sequence and flanking BsaI restriction sites, making it compatible with GreenGate-based CRISPR vectors. The PCR amplicons were purified using the E.Z.N.A.^®^ Gel Extraction Kit (Omega Bio-tek) and cloned into BsaI-compatible sites of the vector CF588 via the GreenGate reaction.

All constructs were transformed and propagated in DH5α *Escherichia coli* cells. Single colonies were selected on solid LB media supplemented with the appropriate antibiotic, propagated in liquid LB media, and plasmids were purified using the E.Z.N.A.^®^ Plasmid DNA Mini Kit I (Omega Bio-tek). All plasmids generated in this study were verified by Sanger sequencing using insert-specific primers (Supplementary Table S4) and are listed in Supplementary Tables S2 and S3).

### RNA extraction and RT-qPCR analysis

For RT-qPCR plants were grown on ½ MS media plates with 1.6% agar at 23°C for 5 days and then transferred to the temperature of interest (10°C, 16°C, 23°C and 27°C) for one more day of growth. Approximately 10 seedlings per biological replicate were collected in Eppendorf tubes^®^, immediately flash-frozen in liquid nitrogen, and stored at −80°C. The frozen plant material was ground to fine powder using a Mixer Mill MM 400 (Retsch) and stainless-steel beads. All steps were performed with cooled-down equipment to prevent premature thawing of the samples. Extraction of RNA was performed using the RNeasy^®^ Plant Mini Kit (Qiagen) following manufacturer’s protocol. RNA concentration and quality were measured using a NanoDrop™ 2000 Spectrophotometer (Thermo Scientific) device and RNA was stored at −80°C till further use.

Reverse transcription was carried out from 1 μg of purified RNA using the RevertAid RT kit (Thermo Scientific) with oligo(dT)_18_ primer. Single strand cDNA was stored at −20°C. RT-qPCR reactions were prepared using the LightCycler® 480 SYBR Green I Master mix. All RT-qPCR assays were performed on CFX™ Real-Time PCR Detection System (Bio-Rad, Inc., Hercules, CA, USA) instruments equipped with either 96-or 384-well blocks using the following program: 5 min at 95°C, followed by 40 cycles of 15 sec at 95°C (denaturation), 30 sec at 55°C (annealing), 20 sec at 72°C (elongation), followed by a melting curve analysis from 55°C to 95°C using 1°C increments. All RT-qPCR assays were performed using 3 biological replicates with 3 technical repetitions each with *TUB2* (AT5G62690) as a reference gene. All primers used for RT-qPCR analyses can be found in Supplementary Table S6.

### Image processing

Figures were processed and prepared for publication using Affinity Photo v1.10.6 and Affinity Designer v1.10.6 (Serif Ltd, https://affinity.serif.com/en-us/).

### Data analysis

Root length was calculated using Fiji 2.14.0 (ImageJ) (Schindelin et al. 2012).

Calculations of the average root length, one-way ANOVA with Tukey’s test to estimate the difference between genotypes, and data plotting were performed using GraphPad Prism version 10.2.2 for Windows, GraphPad Software, Boston, Massachusetts USA, www.graphpad.com. Similarly, calculation of the average seed abortion rate, Tukey test to eliminate the outliers, and data plotting were performed using GraphPad Prism.

## Acknowledgements

We would like to thank the group of Prof. Wolf L. Eiserhardt from Aarhus University for support in phylogenetic analysis and GenomeDK cluster for providing computational resources to perform necessary calculations and support that contributed to these research results. Prof. Christian Fankhauser, Lausanne University, Lausanne, Switzerland, for providing the CF588 vector. Dmitry Malyshev for help with language editing. This research was funded by grants from the Knut och Alice Wallenberg Stiftelse (KAW 2018.0202) and FORMAS (2023-01077) to M.S.

## Author contributions

V.D. performed most experiments and analyzed the data. N.R.M generated and analyzed most of the intron deletion lines. R.M.B. generated and analyzed the *PCPL*/*SmE2* CRISPR/Cas9 mutants. W. L. E. provided guidance on the phylogenetic part of the research. V.D. and M.S. wrote the manuscript with input from all the authors.

## Data availability

Data supporting this study’s findings, which are not already included in the supplementary material, are available from the corresponding author upon reasonable request.

## Supplementary information

**Supplementary Method S1.** Command line instructions used for phylogenetic analysis.

**Supplementary Fig. S1.** *Arabidopsis thaliana* SmE protein structure.

**Supplementary Fig. S2.** CRISPR/Cas9-generated PCPL mutant alleles

**Supplementary Fig. S3.** Plant phenotypes at 27°C and 10°C.

**Supplementary Fig. S4.** Plant phenotypes at 23°C and 16°C.

**Supplementary Table S1.** *SmE* genes used in phylogenetic analysis.

**Supplementary Table S2.** GreenGate plant binary plasmids used in this study.

**Supplementary Table S3.** Summary of plasmids cloned by overlap PCR used in this study.

**Supplementary Table S4.** Oligonucleotides used in this study.

**Supplementary Table S5.** Summary of GreenGate entry modules generated in this study, including information on PCR primers used for amplification.

**Supplementary Table S6.** Primers used for RT-qPCR.

